# Phenotypic and microsite plant individual variation determine the pollination network structure and its functional consequences

**DOI:** 10.1101/2020.03.03.974899

**Authors:** Blanca Arroyo-Correa, Ignasi Bartomeus, Pedro Jordano

## Abstract

The biotic and abiotic context of individual plants in animal-pollinated plant populations may influence pollinator foraging behaviour and therefore how the pollen flow occurs. Thus, this variation among conspecifics within a given plant population can ultimately influence the plant reproductive success. Here we used a fine-scale, well characterized population of *Halimium halimifolium* in combination with exponential random graph models (ERGMs) to assess how the intrapopulation variation in intrinsic (i.e. phenotype and phenology) and extrinsic (i.e. microsite) plant attributes configures individual plant-pollinator networks and its functional consequences. We found that pollinator visitation patterns and the emerging network configuration were associated with both intrinsic and extrinsic plant attributes, such as the number of flowers, the flowering synchrony and the cover of intraspecific and interspecific neighbours. Both intrinsic and extrinsic plant attributes also affected the plant female reproductive success directly and indirectly - through its effects on the probability of conspecifics plants to share pollinators. Our study opens up new approaches to assess and predict the functional consequences of context-dependency in plant-pollinator interactions, especially under global change scenarios, where the ecological context of individual plants is likely to change.

## Introduction

In animal-pollinated plant populations, the foraging movements of pollinators determine how pollen transfer occurs among individuals and the resulting reproductive output (Morris et al. 1995). Within a generalist pollination system, a plant population is often composed of individuals differing in their level of attraction to pollinators, and thus, the realized generalization of each individual is often a subset of its potential (e.g. Herrera 2005, Gómez et al. 2007). As a consequence, we would expect the pattern of pollen transfer to be structured and non-random (Valverde 2017). These mating patterns generated by the establishment of complex individual plant-pollinator interactions importantly influence functional aspects of the mating system by determining population dynamics and genetic diversity patterns.

Individual plants within most populations are phenotypically diverse and differences in floral and vegetative traits (e.g. quality of rewards, plant and flower size or flowering phenology) among plants may determine differences in their pollination niche due to distinctive preferences by different pollinator guilds (Faegri & van der Pijl 1979, Gómez et al. 2013). Hence, individual plants displaying similar phenotypic traits are expected to interact with a similar pollinator assemblage, promoting mating events among those plants (Gómez et al. 2011). Plant populations are also usually composed of individuals with different phenologies (i.e. flowering schedules) (Primack 1985). Likewise, dissimilarities in phenological patterns among pollinator species may create temporal variation in pollinator availability (Cane et al. 2005), which might lead to differences in the pollinator assemblage interacting with those plants diverging in flowering phenology (Fox 2003). Therefore, the flowering synchrony among conspecifics can determine the level of pollinator sharing and thus, the intrapopulational assortativity in mating (Weis et al. 2005).

The variation in the pollination niche among individual plants may also be influenced by the spatial position of those plants within the population (Dupont et al. 2014, Rodríguez-Rodríguez et al. 2017). The spatial position determines the ecological context of individual plants, such as the microtopographical conditions and the local plant community to which they are exposed (e.g., intraspecific and interspecific competition for resources). This variation in microhabitat among plants may generate different aspects of context-dependency of interaction outcomes (Chamberlain et al. 2014). The floral display at the neighbourhood level directly affects the foraging patterns of interacting animals within a plant population (Ghazoul 2005), potentially influencing the mating events among plants (Nattero et al. 2011, Rodríguez-Rodríguez et al. 2015). However, most plant-pollinator interaction studies do not address the spatial component, which might entail important constraints considering the fine-scale division of resources among insect pollinators within a plant population (Dupont et al. 2014) and the spatial heterogeneity in neighbourhood composition (Janovský et al. 2013).

Plant-pollinator communities have been widely studied during the last decades using complex bipartite interaction networks (Bascompte & Jordano 2007), where pollinator species are connected to the plant species they visit. However, species-based networks overlook the existing intraspecific variation by pooling all the individuals in a single group (Tur et al. 2015). In contrast to the community approach, individual-based networks can be used to unveil processes taking place at the population level (e. g. Gómez et al. 2011, Dáttilo et al. 2014, Valverde et al. 2016, Rodríguez-Rodríguez et al. 2017). For instance, in a bipartite individual-based pollination network within a plant population, pollinator species are connected to individual plants they visit. When it is projected into a unipartite network, individual plants of a population are visualized as nodes and the links connecting those nodes represent the level of pollinator sharing, which may serve as a proxy of mating probabilities (Fortuna et al. 2008, Gómez et al. 2011). Thus, the unipartite network projection of plant interconnectedness via shared pollinators allows to elucidate how interactions and consequently potential pollen transfer events are structured within a plant population.

Intrapopulation variation in phenotypic traits, microsite and flowering schedule can therefore affect the network topological position of individual plants (i.e. the relative position of individual plants within the unipartite pollination network). While there are clear hypotheses suggesting that those plants occupying central positions might benefit from a higher number of connections to conspecifics via shared pollinators, increasing the probability of mating events and hence the reproductive success (Gómez & Perfectti 2011, Dupont et al. 2014), it is not trivial how to capture species network topologies into interaction outcomes. Recently developed exponential random graph models (ERGM) allow to evaluate the influence of individual plant attributes on the configuration of the complex network, advancing network studies from descriptive metrics into a more cohesive predictive framework. Such models are a useful tool to account for the overall structure of networks as a function of attributes of the interacting nodes (Snijders et al. 2010). Although this predictive and modeling framework has been widely used in social network studies in recent years, it has been rarely considered for ecological network analysis (but see Miguel et al. 2018).

Here we aim to investigate how the intrapopulation variation in intrinsic (i.e. phenotypic traits and phenology) and extrinsic (i.e. microsite) plant attributes shapes the structure of individual plant-pollinator networks and the female plant reproductive consequences. To address this question, we recorded interactions between pollinator functional groups and individual plants in a population of the Mediterranean shrub *Halimium halimifolium* (Cistaceae). Specifically, we assess (i) how the pollinator visitation patterns and the emerging configuration of the bipartite individual plant-pollinator network are related to intrinsic and extrinsic plant attributes, and (ii) to what extent the female reproductive success of individual plants is influenced by plant attributes and the structure of the unipartite plant-plant network derived from pollinator sharing among conspecifics.

## Materials & Methods

### Study site

The study was performed in Doñana National Park (37°07’52.2”N 6°31’40.9”W, 12 m a.s.l.) within an area of 1.2 ha bounded on its four sides by a water stream, pine forests and grasslands. Doñana National Park is located in the Atlantic coast of southwestern Spain, in an area with a Mediterranean type climate where the vegetation is composed mainly of sclerophyllous shrublands. The study area is largely dominated by *Halimium halimifolium* (Cistaceae), a mediterranean shrub that grows in the slopes of stabilized sand dunes (Díaz-Barradas & García-Novo 1987). Other insect-pollinated plant species, such as *Halimium commutatum, Lavandula stoechas* and *Cytisus grandiflorus*, are also present in our study area (Fig. 1a, Fig. S1a). However, because *H. halimifolium* was the only insect-pollinated plant species with flowers during this study, we did not consider competition for pollinators and facilitation among the different plant species.

**Fig. 1.**
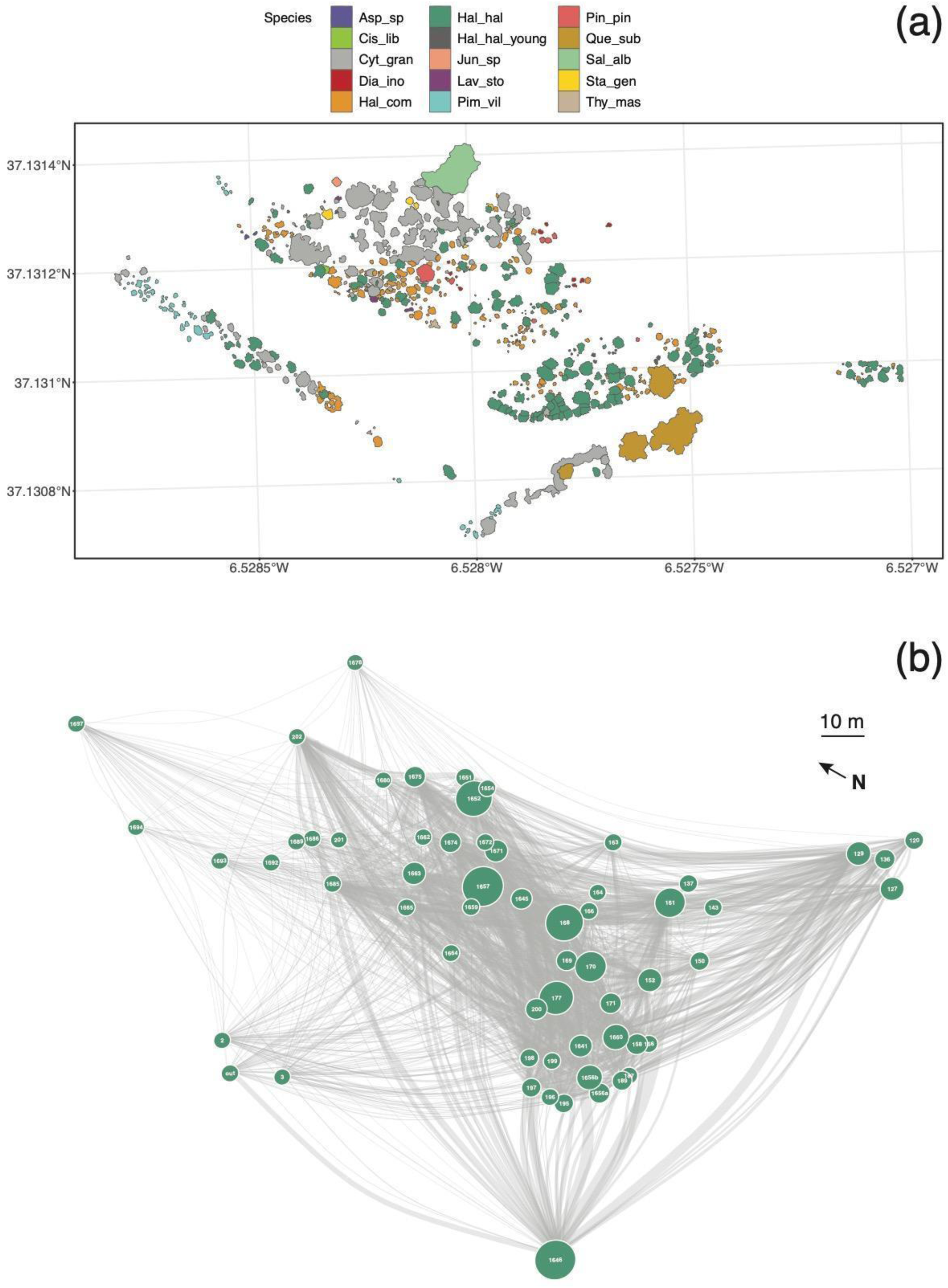
**(a)** Spatial distribution of all individual plants and *Halimium halimifolium* individual plants surveyed in the study site. Each polygon represents an individual plant and the colour refers to the species it belongs to: Asp_sp: *Asparragus* sp., Cis_lib: *Cistus libanotis*, Cyt_gran: *Cytisus grandiflorus*, Dia_ino: *Dianthus inoxianus*, Hal_com: *Halimium commutatum*; Hal_hal: *Halimium halimifolium*, Hal_hal_young: *Halimium halimifolium* saplings, Jun_sp: *Juncus* sp., Lav_sto: *Lavandula stoechas*, Pin_pin: *Pinus pinea*, Que_sub: *Quercus suber*, Sal_alb: *Salix alba*, Thy_mas: *Thymus mastichina*, Sta_gen: *Stauracanthus genistoides*, Pim_vil:: *Pimpinella villosa*. **(b)** Weighted unipartite network illustrating the pattern of shared pollinator functional groups among *H. halimifolium* individual plants. Two individual plants are linked if they shared at least one pollinator functional group visiting their flowers. Link width is proportional to the number of interactions shared between two individual plants. Node size refers to the female reproductive success (FRS) while node position indicates the spatial location (X-Y coordinates) of individual plants.

*Halimium halimifolium* plants have large (up to 62 mm in diameter) hermaphrodite yellow flowers (Fig. S1b-c) that bloom in late-spring (May–June; Herrera 1988). Each individual plant can produce up to one thousand flowers at once and they are pollinated mainly by Coleoptera, Hymenoptera and Diptera (Herrera 1988). Flower opening occurs synchronously each day within the population and each flower lasts exclusively a few hours. Because *H. halimifolium* is predominantly self-incompatible, cross-pollination promoted by insect visitations is necessary to enhance seed production (Talavera et al. 1997).

### Sampling

We conducted surveys to record pollinator visitations in the study area during the peak flowering period of *H. halimifolium* (20 days, May 2019 - June 2019). Prior to data collection, all individual plants of the population (160 individuals) were labelled. Interactions with pollinator species were recorded in 60 of those individuals, which were selected using a stratified random sampling where both isolated and aggregated plants were well represented (Fig. S3). We performed four 30-min surveys in each individual plant using cameras (GoPro Hero7, GoPro Inc., Germany). Therefore, each of the 60 labelled plants was surveyed for 120 min and hence, the entire survey lasted 7200 min. To avoid within-day temporal bias in pollinator assemblage, these surveys per plant were carried out in different times during the time period with open flowers (8:30 to 13:30). This sampling period encompasses the time of the day where maximum activity of flower visitors to *H. halimifolium* occurs (Talavera et al. 1997). We assessed the completeness of sampling effort calculating Chao asymptotic estimators (see Appendix S1, Fig. S2).

During each survey interval, we recorded the number of open flowers in the individual plant, the identity of pollinator species visiting the flowers and the number of times they interacted with the flowers (i.e. frequency of interactions). Pollinator species were considered as all those insects landing on the flower and touching its reproductive structures. We classified pollinator species into eight functional groups that were defined considering similarities in feeding habits, foraging behavior and body size (Table S1). A single representative specimen of each pollinator species was captured and identified.

We constructed a bipartite weighted network linking each individual plant with the pollinator functional groups visiting its flowers. The individual-based network was built by creating an adjacency matrix *A*, where elements *a*_ij_= number of interactions between the pollinator functional group *i* and the individual plant *j*, and zero otherwise.

To depict the pattern of shared pollinators among individual plants, we generated the unipartite projection of matrix *A* by calculating the number of interactions shared by any two individual plants. The unipartite projection of matrix *A* for the *P* plants would be: *A*_P_= *AA*_T_, where *A*_T_ is the transpose of *A*. In this unipartite network, two plants are linked if they share at least one interaction with the same pollinator functional group. Hence, individual plants are not connected by plant-to-plant movements of individual pollinators, but shared functional groups of pollinators. We assumed that sharing pollinator functional groups is a surrogate indicator of pollen transfer among individual plants given that most pollinators within a functional group have similar foraging behavior (Fenster et al. 2004, Gómez & Perfectti 2011). Thus, pollinator individuals belonging to the same functional group are likely to show the same interaction patterns with individual plants. Two plants that are connected in the unipartite network, and thus shared pollinator groups, will be more likely to be involved in mutual pollen transfer, and a potential mating event, than two plants with very different sets of pollinators. We used the link weights estimated in this way as a proxy for probability of mating events mediated by a given pollinator functional group (see Gómez et al. 2011, Rodríguez-Rodríguez et al. 2017).

### Plant attributes

We obtained a set of intrinsic and extrinsic attributes for each surveyed individual plant (Appendix S1), but we exclusively retained in the analysis those which were not redundant after testing for multicollinearity using VIF factors with a threshold of 2 (‘VIF’ R package; Lin 2012, Table S2). We included several microsite characteristics as extrinsic variables related to competition and microtopographical conditions: (i) cover of intraspecific and interspecific neighbours within 1m and 2m radius, (ii) minimum distance to the stream, (iii) minimum distance to the habitat edge and (iv) distance to the nearest tree. The intrinsic variables of each individual plant included in the analyses comprised the following phenotypic traits: (i) maximum height (m), (ii) total number of flowers per individual during the peak flowering season, (iii) maximum diameter of the corolla (mm) and (iv) size of the flower guides (mm). To estimate individual overlap in flowering time within the entire population, we calculated a flowering synchronization level by modifying the index proposed by Marquis (1988) (Appendix S1). All extrinsic variables as well as the maximum plant height were estimated using aerial images, an orthomosaic and a 3D surface model obtained with drone flights (see Appendix S1).

### Female fitness estimation

To quantify the female reproductive success (FRS hereafter) of each individual plant, we first calculated the fruit set as the proportion of flowers setting fruit. We collected 10 inflorescences per plant and counted the total number of flower buds produced and the number of fruits. Second, we obtained the seed production per fruit in each plant by sampling 5 fully-developed fruits per plant to count the number of seeds and we measured the seed mass using a precision scale. An overall female reproductive success (FRS) for each plant was estimated by weighing the total number of seeds per plant by seed mass (Appendix S1).

### Topological position of individual plants

The relative contribution of individual plants within the pollination network was calculated in terms of their centrality (Freeman 1978), which indicates how well connected is a given individual plant with the rest of the co-occurring conspecifics via shared pollinator functional groups. We assumed that an increased fraction of shared pollinators reveals a higher potential for pollen transfer/reception among two individuals relative to other individual pairs with different pollinators. We estimated two measures of centrality from the unipartite projection of the weighted network: (i) closeness centrality, which indicates how well pollen originating from a focal plant can reach other plants or how well pollen from conspecifics can reach this focal plant, and (ii) betweenness centrality, which represents how well an individual plant facilitate the exchange of pollen among all nodes within the network (see Appendix S1 for more detailed definitions of these metrics). Centrality metrics were calculated with the ‘bipartite’ R package (Dormann et al. 2008).

### Statistical analyses

#### Association between plant attributes and interaction patterns with pollinators

The relationship between pollinator visitation patterns and plant attributes was tested with a Canonical Correlation Analysis (CCA; Borcard et al. 2018) by comparing two multivariate datasets, one including pollinator visitations to plants and the other one including intrinsic and extrinsic plant attributes. CCA determines a set of canonical variates, which are linear combinations of the variables within each dataset that best explain the variability both within and between sets. The extent to which the full model captures the relationship between two datasets is determined by the canonical correlation coefficient (R^2^) between canonical variates, which reflects the total variance from datasets explained by the variates. The significance of the canonical correlation was assessed with a Wilk’s lambda test (Nimon et al. 2010).

#### Ecological correlates of individual plant-pollinator network structure

To analyze the ecological variables producing the overall structure of the weighted individual-based pollination network emerging from interaction patterns we built exponential random graph models (ERGMs) (Lusher et al. 2013). The design of ERGMs is analogous to the classical generalized linear models (GLMs) and implements a Markov chain Monte Carlo maximum likelihood to calculate parameter estimates. ERGMs test hypotheses about underlying mechanisms shaping networks by modeling how the likelihood of edge formation (i.e. the presence of interaction links between a pollinator functional group and an individual plant) is affected by specific variables. Therefore, the presence or absence of network links and their configuration is defined as the response variable, while the predictor variables can be different kinds of network structures (i.e. endogenous variables that specify different configurations, defined to be sets of possible edges among a subset of the nodes in the network) or node and edge attributes (i. e. exogenous variables, such as plant phenotypic traits or pollinator characteristics). The simplest ERGM controls for endogenous effects originated just from the pattern of links among nodes and therefore this model only incorporates statistics as functions of the network structure (Morris et al. 2008). However, the probability of a link between two nodes is expected to depend not only on the presence or absence of links joining two nodes, but also on exogenous effects derived from the nodes’ attributes. We can incorporate these effects into the ERGM including measured attributes of the individual plant nodes as additional statistics in the exponential term.

In our random graph models, we considered phenotypic traits and microsite characteristics of individual plants as predictor variables and the structure of the weighted bipartite plant-pollinator network as the response variable. Thus, our aim was to assess the contribution of each predictor variable to the overall network configuration by associating the structure of the plant-pollinator network to specific attributes of plant nodes. To determine which ecological variables are the main contributors to the network structure, we set four models including different predictors (Appendix S1). All predictor variables were previously scaled to allow meaningful comparisons among those variables.

#### Fitness consequences

To analyze the reproductive consequences of plant-pollinator interactions, we performed an additional ERGM to assess the association between the overall patterns of interactions among individual plants derived from shared pollinator functional groups and FRS. By doing so, we aimed to investigate whether higher final FRS of two linked plants was related to the probability of having shared pollinators. We might expect such an effect whenever highly effective pollinator functional groups selectively visit a subset of plants and this results in higher overall FRS. In this model, we included the structure of the unipartite plant-plant network as the response variable and FRS, phenotypic traits and microsite as predictor variables. As a consequence of optimal foraging strategies of pollinators, they tend to move at short distances and thus, closer plants are more likely to share pollinators than further plants. Therefore, we may expect that the spatial distance between individual plants would affect the level of pollinator sharing. To account for this spatial effect, the ERGM with the unipartite network as the response variable allows as to include a covariance matrix with spatial distance between individual plants as a predictor variable.

The direct effects (i.e. not mediated by pollinator sharing) of plant attributes on the female reproductive success was tested using a generalized linear models (GLM) with quasibinomial errors. This model included the normalized FRS as the response variable and plant attributes as predictor variables. We also included centrality metrics as predictor variables to account for the effect of the topological position of plants in the unipartite network on FRS. Prior to model fitting, we tested for spatial autocorrelation of FRS and intrinsic plant attributes estimating Moran’s I with the ‘ape’ R package (Paradis & Schliep 2018). All predictor variables were previously scaled to allow meaningful comparisons among those variables. The contribution of the different fitness components to FRS was assessed using the ‘relaimpo’ package in R (Grömping 2006). This package fits linear models with FRS as the response variable and its components (i.e. total number of flowers, fruit set, number of seeds per fruit and seed mass) as predictor variables and estimates the R^2^ contribution of those components to FRS averaged over resampled orderings among regressors.

ERGMs were computed using the ‘sand’ and ‘ergm’ R packages (Hunter et al. 2008, Handcock et al. 2018, Kolaczyk & Csárdi 2017). The Canonical Correlation Analysis was performed using the ‘CCA’ R package (González & Déjean 2012). All analyses and graphical representations were performed using R software version 3.5.3 (R Core Team 2019).

## Results

We analyzed 3600 minutes of video recordings in which we registered 2778 interactions between 60 individual plants and 22 pollinator species that were classified into eight functional groups (Table S1). Along the study population, most pollinator visits were performed by small bees (42.09%), medium size bees (33.87%) and large beetles (11.71%). We observed substantial variation among individual plants in both extrinsic (i.e. microsite characteristics) and intrinsic (i.e. phenotypic traits) attributes (Table S2).

### Association between plant attributes and interaction patterns with pollinators

We found a significant association between pollinator visitation to individual plants and plant attributes (Table S3). The first canonical variate was the only one significant and explained 72% of the total variance between the two original datasets (Wilk’s λ= 0.047, F= 1.67; df= 96, 279.72; p< 0.001). This canonical variate was highly correlated with maximum height of the plant (−0.333), total number of flowers during the flowering peak (0.828), distance to the nearest tree (0.223) and to the habitat edge (−0.229), cover of intraspecific neighbours in r= 1m (0.221) and r= 2m (−0.397); and cover of interspecific neighbours in r= 1m (−0.274) and r= 2m (0.582). Regarding pollinator functional groups, the first canonical variable was mostly correlated with large bees (0.359), large beetles (0.572) and medium size bees (0.591).

### Ecological correlates of individual plant-pollinator network structure

The number of edges in the bipartite network (i.e. unique links between individual plants and pollinator functional groups) was *L(y)*= 208 (Fig. S4). We did not find an edge effect in the ERGM fitted with the bipartite network as the response variable (Table 1), suggesting that the probability of observing the given network configuration was not influenced by the number of interactions and its distribution among nodes. This means that the observed data significantly fit a Bernoulli random graph model where, for a given pair of nodes, the presence or absence of an edge between that pair is independent of the status of possible edges between any other pairs of nodes. Therefore, additional variables (exogenous) are needed to explain the observed structure of the plant-pollinator network. The full ERGM with plant phenotypic traits and microsite characteristics as predictor variables was the one that better explained the network structure (Table S4).

**Table 1.**
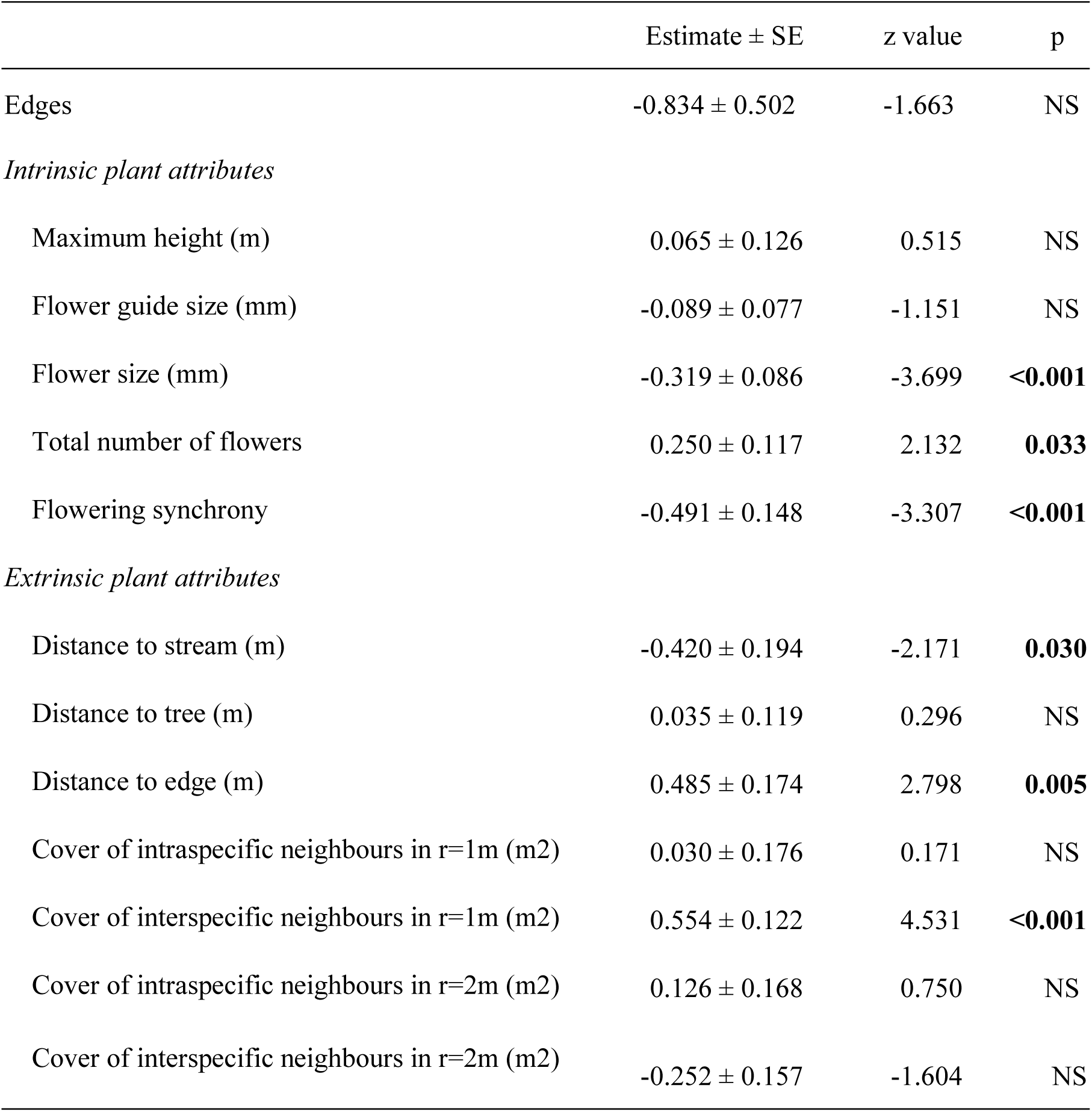
Summary of the exponential random graph model (ERGM) showing the effects of individual plant attributes in the bipartite pollination network structure. The edges effect assesses the influence of the interaction patterns among individual plants and pollinator functional groups on the configuration of the bipartite network. Variables with significant effects indicates an increase or decrease in the probability of an individual plant to interact with the set of pollinator functional groups. Significant values (p< 0.05) appear in boldface.

| | Estimate ± SE | z value | p |
| --- | --- | --- | --- |
| Edges | -0.834 ± 0.502 | -1.663 | NS |
| *Intrinsic plant attributes* |  |  |  |
| Maximum height (m) | 0.065 ± 0.126 | 0.515 | NS |
| Flower guide size (mm) | -0.089 ± 0.077 | -1.151 | NS |
| Flower size (mm) | -0.319 ± 0.086 | -3.699 | **<0.001** |
| Total number of flowers | 0.250 ± 0.117 | 2.132 | **0.033** |
| Flowering synchrony | -0.491 ± 0.148 | -3.307 | **<0.001** |
| *Extrinsic plant attributes* |  |  |  |
| Distance to stream (m) | -0.420 ± 0.194 | -2.171 | **0.030** |
| Distance to tree (m) | 0.035 ± 0.119 | 0.296 | NS |
| Distance to edge (m) | 0.485 ± 0.174 | 2.798 | **0.005** |
| Cover of intraspecific neighbours in r=1m (m2) | 0.030 ± 0.176 | 0.171 | NS |
| Cover of interspecific neighbours in r=1m (m2) | 0.554 ± 0.122 | 4.531 | **<0.001** |
| Cover of intraspecific neighbours in r=2m (m2) | 0.126 ± 0.168 | 0.750 | NS |
| Cover of interspecific neighbours in r=2m (m2) | -0.252 ± 0.157 | -1.604 | NS |

In any ERGM, the model coefficients for exogeneous variables (Table 1) represent the effects of attributes of the nodes on the likelihood of any pair of plant-pollinator group establishing a link via an interaction, as expressed from the bipartite network described above. They are interpreted as a conditional log-odds ratio for visitation (links) for any pair of plant-pollinator interaction increasing (or decreasing) as a function of a given change in the specific attribute. For example, how increasing flowering synchrony increases the odds of visitation by more different functional groups of pollinators. Thus, our model showed that increasing the total number of flowers in individual plants increased the odds of being visited by a greater number of distinct functional groups of pollinators by a factor of exp(0.250)= 1.284, or nearly 30% (Table 1). Additional plant attributes that increased the odds of more diversified plant-pollinator interactions were the cover of interspecific neighbours in a 1m radius and distance to the habitat edge. The odds of these interactions decreased with flower size, flowering synchrony and distance to stream. For most significant exogeneous variables, the coefficient differed from zero by at least one standard error, suggesting a sizeable effect of these variables on the likelihood of plant-pollinator link establishment.

### Fitness consequences

The number of edges in the unipartite projection of the individual plant-pollinator network (i.e. links among individual plants sharing pollinator functional groups) was *L(y)*= 1478 (Fig. 1b). This unipartite projection represents the way individual plants share pollinator groups and hence a potential for mating events mediated by these functional groups acting as pollinators. The edge effect was highly significant in the ERGM fitted with the unipartite network as the response variable, i.e. the odds of a link between two individual plants was proportional to the degree of individual plants (Table 2). Therefore, the presence or absence of an edge between a given pair of individual plants depended on the status of possible edges between any other pairs of plants in the unipartite network. We found a significant positive association between the probability of two individual plants to share pollinators and FRS (exp(2.837)=17.064). Variables that increased the odds of sharing pollinators between individual plants were the height of the plant, flower size, distance to the stream, distance to the nearest tree and cover of intraspecific neighbours in a 2 m radius. Meanwhile, the odds of two individual plants to share pollinators decreased with the spatial distance between plants, flowering synchrony, distance to habitat edge and cover of intraspecific neighbours in a 1 m radius (Table 2). Again, for most significant exogeneous variables, the coefficient differed from zero by at least one standard error, suggesting a sizeable effect of these variables on the likelihood of link establishment among individual plants.

**Table 2.**
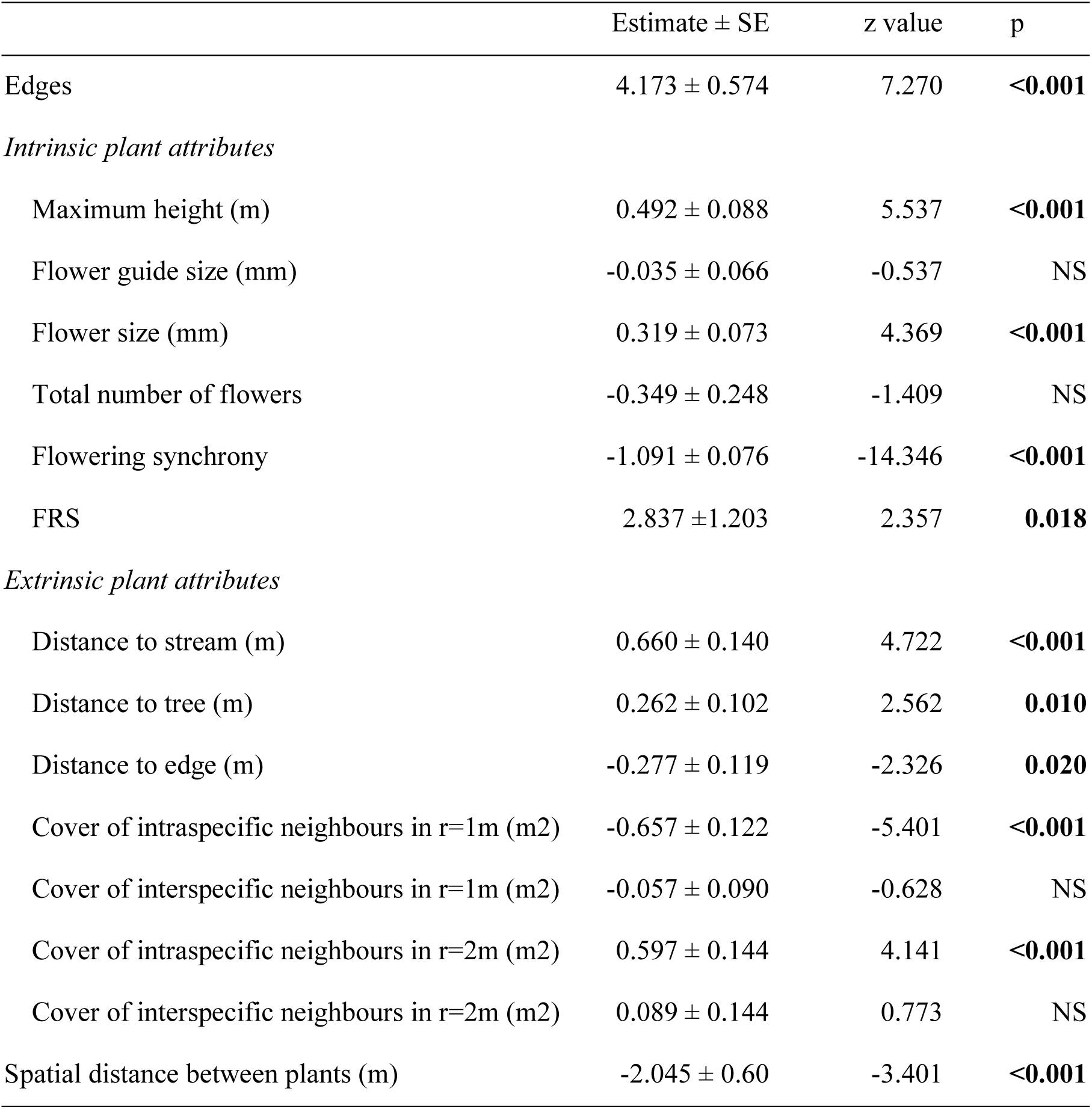
Summary of the exponential random graph model (ERGM) evaluating the association between individual plant attributes, female reproductive success (FRS) and the structure of the unipartite network constructed from interactions among plants sharing pollinator functional groups. The edges effect assesses the influence of interaction patterns among individual plants on the configuration of the unipartite network. Variables with significant effects indicates an increase or decrease in the probability of two individual plants to interact (i.e. to share pollinator functional groups). Significant values (p< 0.05) appear in boldface.

| | Estimate ± SE | z value | p |
| --- | --- | --- | --- |
| Edges | 4.173 ± 0.574 | 7.270 | **<0.001** |
| *Intrinsic plant attributes* |  |  |  |
| Maximum height (m) | 0.492 ± 0.088 | 5.537 | **<0.001** |
| Flower guide size (mm) | -0.035 ± 0.066 | -0.537 | NS |
| Flower size (mm) | 0.319 ± 0.073 | 4.369 | **<0.001** |
| Total number of flowers | -0.349 ± 0.248 | -1.409 | NS |
| Flowering synchrony | -1.091 ± 0.076 | -14.346 | **<0.001** |
| FRS | 2.837 ± 1.203 | 2.357 | **0.018** |
| *Extrinsic plant attributes* |  |  |  |
| Distance to stream (m) | 0.660 ± 0.140 | 4.722 | **<0.001** |
| Distance to tree (m) | 0.262 ± 0.102 | 2.562 | **0.010** |
| Distance to edge (m) | -0.277 ± 0.119 | -2.326 | **0.020** |
| Cover of intraspecific neighbours in r=1m (m<sup>2</sup>) | -0.657 ± 0.122 | -5.401 | **<0.001** |
| Cover of interspecific neighbours in r=1m (m<sup>2</sup>) | -0.057 ± 0.090 | -0.628 | NS |
| Cover of intraspecific neighbours in r=2m (m<sup>2</sup>) | 0.597 ± 0.144 | 4.141 | **<0.001** |
| Cover of interspecific neighbours in r=2m (m<sup>2</sup>) | 0.089 ± 0.144 | 0.773 | NS |
| Spatial distance between plants (m) | -2.045 ± 0.60 | -3.401 | **<0.001** |

Female reproductive success (FRS) of individual plants and intrinsic plant attributes were not spatially autocorrelated (Table S5). The GLM revealed that FRS was significantly associated with both the topological position of individual plants within the inferred mating network and plant attributes (variation explained= 87.29%; Table 3). We found that FRS increased with closeness centrality, total number of flowers during the flowering peak and flowering synchrony (Fig. 2); and decreased with distance to the stream and cover of intraspecific neighbours in a 2m radius (Table 3). Differences in FRS were mainly determined by the total number of flowers during the flowering peak (86.10% of the effects) and the fruit set (3.97% of the effects) (Table S6).

**Table 3.** Results of the generalized linear model (GLM) evaluating the effects of the topological position and individual plant attributes on female reproductive success (FRS). The fitted GLM accounted for 87.29% of the variation. FRS was estimated as the total number of seeds per individual weighted by the seed mass and normalized (See *Materials & Methods: Female fitness estimation*). Significant values (p< 0.05) appear in bold.

| | Estimate ± SE | t value | p |
| --- | --- | --- | --- |
| (Intercept) | -2.561 ± 0.131 | -19.587 | **<0.001** |
| *Topological position* |  |  |  |
| Betweenness | 0.168 ± 0.158 | 1.060 | NS |
| Closeness | 0.623 ± 0.263 | 2.367 | **0.022** |
| *Intrinsic plant attributes* |  |  |  |
| Maximum height (m) | -0.318 ± 0.167 | -1.905 | 0.063 |
| Flower guide size (mm) | 0.076 ± 0.121 | 0.632 | NS |
| Flower size (mm) | -0.113 ± 0.138 | -0.817 | NS |
| Total number of flowers | 1.497 ± 0.166 | 9.016 | **<0.001** |
| Flowering synchrony | 0.714 ± 0.265 | 2.694 | **0.010** |
| *Extrinsic plant attributes* |  |  |  |
| Distance to stream (m) | -0.469 ± 0.214 | -2.191 | **0.034** |
| Distance to tree (m) | -0.238 ± 0.175 | -1.361 | NS |
| Distance to edge (m) | 0.003 ± 0.225 | 0.013 | NS |
| Cover of intraspecific neighbours in r=1m (m2) | 0.256 ± 0.22 | 1.164 | NS |
| Cover of interspecific neighbours in r=1m (m2) | -0.184 ± 0.221 | -0.833 | NS |
| Cover of intraspecific neighbours in r=2m (m2) | -0.424 ± 0.199 | -2.132 | **0.038** |
| Cover of interspecific neighbours in r=2m (m2) | -0.024 ± 0.21 | -0.115 | NS |

**Fig. 2.**
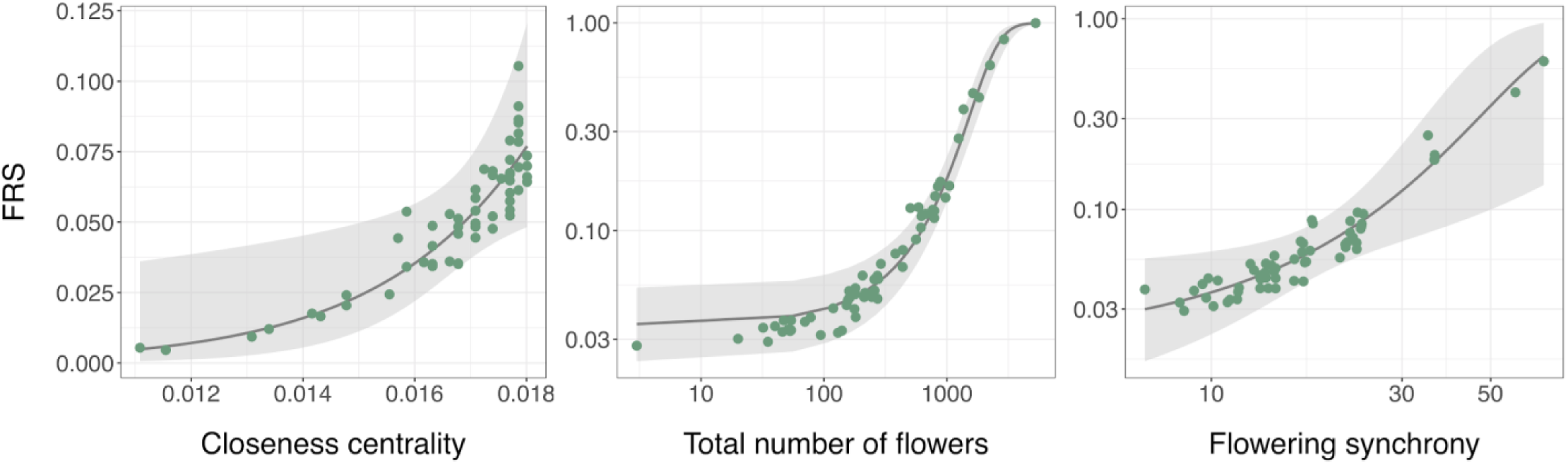
Partial residual plots illustrating the variability of female reproductive success (FRS) of individual plants explained by closeness centrality, total number of flowers and flowering synchrony, when controlling for the effects of the remaining variables included in the model. These three attributes of individual plants were the predictor variables with higher effects on FRS. Points represent individual plants surveyed in the studied population and the grey area indicates the 95% confidence interval for the fitted values. Axis scale differ among plots to enhance the visualization of the data. Note log scale on both axes for the plots showing the effects of the total number of flowers and flowering synchrony.

## Discussion

Our findings revealed how intrinsic and extrinsic attributes of individual plants can influence the establishment of complex individual plant-pollinator interactions and their consequences for individual reproductive outcomes. We found that both the visitation patterns of pollinator functional groups to individual plants and the overall network configuration were associated with both intrinsic (i.e. phenotypic traits and flowering phenology) and extrinsic (i.e. microsite) plant attributes. We also found that those plant attributes affected the female reproductive success of individual plants directly and indirectly - through their effects on the topological role in the plant-pollinator network and the probability of those individual plants to share pollinators with conspecifics. Overall, these results suggest that the ability of individual plants to maintain the cohesion of the entire population (i.e. the pollen transfer via mutualistic interactions) may ultimately be influenced by their phenotypic traits and ecological context within the population.

### Association between plant attributes and interaction patterns with pollinators

The establishment of mutualistic interactions is influenced by both the abiotic and biotic context in which it takes place and therefore it is highly dynamic over space (Chamberlain et al. 2014). We found that the patterns of pollinator visitation to individual plants were mainly associated to both intrinsic (plant attributes related with plant height and flower abundance) and extrinsic (microsite characteristics) attributes of plants. Thus, our findings support the notion that pollination visitation patterns are significantly influenced by extrinsic attributes besides plant phenotypic traits. This result ties well with other studies wherein the local microclimatic variation (e.g., solar irradiance, humidity and temperature) influenced pollinator composition, activity and behavior at flowers (Herrera 1995, Potts et al. 2005). Our results expand these findings by suggesting these influences extend to the overall structure of the pollination network at the population level (see below), potentially influencing mating patterns and local gene flow.

Similar to previous studies (Gómez & Perfectti 2011, Weber & Kolb 2013, Dupont et al. 2014) we found that intraspecific trait variation in flower abundance and height among plants also explained differences in the pollinator assemblage visiting individual plants. However, contrary to these studies we did not find a significant effect of floral phenotype (i.e. flower size and floral guide size) on the visitation patterns. This might respond to the fact that the floral phenotype of *H. halimifolium* in our population did not show wide variation as to promote differential attraction for pollinators (Van der Niet et al. 2014). Alternatively, if pollinators visiting *H. halimifolium* plants have a highly generalist foraging behaviour, the observed variation in floral phenotype may not have a strong impact on pollinator attraction and therefore on pollinator visitation patterns. Another possible interpretation is that the variation in other intrinsic attributes, such as plant height, or extrinsic attributes could be more important for pollinator attraction that differences in floral phenotypes.

### Ecological correlates of individual plant-pollinator network structure

By using exponential random graph models (ERGMs) we were able to further recognize specific plant attributes at different scales (individual phenotypic traits and microsite) that shape the observed individual plant-pollinator network. ERG models allow the exploration of plant attributes directly affecting the composition and generalization level of individual flower visitor assemblages. This in turn allows us to establish an effective bridge between variation in interaction network topologies and fitness variation in natural populations. We found that a set of extrinsic and intrinsic ecological variables explained the odds individual plants have in establishing interactions with specific pollinator functional groups in our study population. Variation in traits of both plant and animal partners are increasingly recognized as forces in structuring ecological networks at the level of interactions among species (Stang et al. 2009, Olesen et al. 2010). Similarly, intraspecific variation in traits can also play an important role in shaping individual-based networks, yet this aspect remains virtually unexplored in the literature. Our findings revealed that the number of flowers produced per individual plant increased the probability of interactions between pollinators and individual plants, increasing the odds for individual plants to be visited by a larger and more diversified array of pollinator functional groups. On the other side, we found that flower size had the opposite effect, decreasing the odds of individual plant-pollinator interactions and resulting in reduced pollinator assemblages.

Contrary to our expectations, the flowering synchrony of individual plants decreased the probability of interacting with pollinators. This result may respond to the fact that a high abundance of flowers within the population can result in the use of resources from multiple individual plants by pollinators, decreasing the chance of a pollinator to visit a given plant individual. However, as previously indicated, we found that the flowering synchrony of individual plants also increased the female reproductive success (FRS). Together, our findings suggest that plants with a higher flowering synchrony, despite having less visits, may benefit from long-distance pollen dispersal and therefore they may be more likely to receive better quality pollen than asynchronous individual plants (see e.g. Castilla et al. 2017).

Although the effect of microsite, especially neighbourhood composition and density, have been previously considered for plant-frugivore (e.g. Carlo 2005, Morales et al. 2012) and plant-pollinator interactions (e.g. Lázaro et al. 2009, Hegland 2014), it has been rarely included in network studies. However, our results demonstrated that the microsite of individual plants was an important driver of the establishment of interactions with pollinators. In light of these results, the context-dependence previously found in species-level interaction networks (Poisot et al. 2015) appears as an important driver of individual-based pollination networks, with direct consequences for fitness variation.

### Fitness consequences

Assessing the consequences of plant-pollinator interaction patterns in the reproductive success of individual plants is crucial to understand the processes taking place at the population level (Wilson & Thomson 1991, Mitchell et al. 2009). As expected, we found that those individual plants with higher closeness centrality in the network showed a higher FRS. Thus, plants sharing pollinators with most other conspecifics may have higher probabilities of pollen exchange and therefore, enhance their reproductive success. However, the variation in FRS was also directly driven by differences in plant attributes related to flower abundance and microsite variables beyond the effects of the topological position of individual plants.

Because we estimated female reproductive success of individual plants as the total number of seeds produced per plant, we expected the total number of flowers per plant to have a significant direct effect on fitness. Moreover, we also found that additional plant attributes affected the female reproductive success. Our results showed that the combination of the topological position in the network and both the intrinsic and extrinsic attributes of individual plants is accurate at predicting the reproductive consequences of plant-pollinator interactions. By contrast, previous work found that the topological role of plants had a higher effect than plant attributes on plant fitness (Gómez & Perfectti 2011, Rodríguez-Rodríguez et al. 2017), but this could be strongly dependent on the spatial and taxonomic (phylogenetic) context. As we did not find a significant association between floral traits, pollinator visitation patterns and FRS, our findings also suggest that floral phenotype in our study population may have limited or absent adaptive value (Herrera 1996). This is likely related to the open, accessible floral design of most Cistaceae species, which may promote visitation by generalist species with limited functional diversification but with adequate effectiveness.

The use of ERG models with the unipartite network derived from pollinator sharing is a useful tool to evaluate how attributes of individual plants influence the odds of sharing pollinators among them and hence, of being involved in reciprocal pollen transfer and mating events. We found that the intrinsic and extrinsic attributes of individual plants significantly influenced the odds of sharing pollinator functional groups. Moreover, our findings revealed that the female reproductive success of individual plants (FRS) was positively associated with the probability of sharing pollinators with conspecifics. Therefore, the odds of sharing pollinators suggest a trend for increasing probabilities of mating among individual plants. Together, these results indicate that the effect of both intrinsic and extrinsic plant attributes on the degree of pollinator sharing among conspecifics can ultimately determine differences in FRS, even at the scale of small populations. The emergence of this pattern of non-random mating events may create a spatial genetic structure within the plant population as a result of the spatially-restricted gene flow generated (Epperson 1993). Thus, our results, bridging the complexity of the pollination network with its direct effects on fitness variation, would contribute to explain the higher levels of genetic structure found in most animal-pollinated plant populations (Vekemans & Hardy 2004). Ultimately, the long-term consequences of spatially-restricted gene flow in our study population would also depend on seed dispersal processes (mostly ant-mediated in *H. halimifolium*) enhancing or erasing the genetic signal generated by the non-random mating events (Fortuna et al. 2008). Future work considering paternal analysis of offspring to infer pollen flow patterns will further our knowledge of how individual plant attributes influence the reproductive outcomes of plant-pollinator interactions.

## Conclusions

Our study opens up new analytical approaches to further understand how complex plant-pollinator networks at the population level are configured. By using individual-based networks and ERGMs we found that both intrinsic and extrinsic plant attributes significantly influenced the overall structure of the individual plant-pollinator network and the reproductive outcomes for individual plants. Hence, our results provide novel insights into the functional consequences of the ecological context of plant–pollinator interactions within a plant population, especially in habitats with patchiness in biotic and abiotic variables that affect pollinator behavior. Our findings may have important implications for the persistence and dynamics of animal-pollinated plant populations under global change scenarios, where the biotic and abiotic context in which individual plants occur are likely to change. Future research should move forward in the application of this modelling approach to assess and predict the potential consequences for population performance of this context-dependency in individual-based pollination networks.

## Acknowledgments

We thank Ricardo Díaz-Delgado for technical support with drone flights and spatial data acquisition. We are grateful to Ana Benítez, Irene Mendoza, Lisieux Fuzessy, Elena Quintero, Jorge Isla and Daniel Pareja for helpful comments. Doñana National Park authorities approved permissions to work in the study area. Financial support was provided by a JAE-Intro fellowship from CSIC (JAEINT18_EX_0080 to BAC) and the Spanish Ministry of Science and Innovation (project CGL2017-082847P to PJ and CGL2017-92436EXP to IB).

