## Appendix for "Phenotypic and microsite plant individual variation determine the pollination network structure and its functional consequences"

This file contains the supplementary tables and figures referred to in the main text.

**Appendix S1**

### 1. Methods

*Completeness of sampling effort*

In order to assess the completeness of sampling effort, we fit the cumulative number of recorded pollinator species and functional groups with increasing number of censuses, calculating the Chao asymptotic estimators with their standard errors using the 'specpool' function in 'vegan' R package (Oksanen et al. 2019; Fig. S2).

*Estimation of extrinsic plant attributes*

In addition to the plant attributes used in the analyses after testing for multicollinearity, we also measured the area (m2), perimeter (m), mean height (m), mean number of flowers per individual during the peak flowering season, and cover of intraspecific and interspecific neighbours within 0.5 m and 3 m radius.

Microsite and spatial location of each individual plant were obtained using aerial images of the study site, with which we generated an orthomosaic together with a 3D digital surface model using the Pix4D software (Pix4D SA, Prilly, Switzerland). Aerial images were acquired with a flight height of 70 m and a pixel size of 1.71 cm using a senseFly SODA photogrammetry camera integrated in an eBee fixed wing UAV (Parrot SA, Paris, France). The information derived from the aerial images was validated using a high-accuracy GNSS handheld (Trimble Geo 7x, Trimble Inc., Wesminster, USA) to register the spatial location and the maximum height of 30 randomly selected plants in the population. We used the orthomosaic to digitize all plants in the study site (n= 546) as polygons and we attributed them to different plant species (Fig. 1a). The polygon layer generated was used to calculate the interspecific and intraspecific neighbourhood area within different size radius and the distance to the stream, habitat edge and nearest tree with the ‘sf’ R package (Pebesma 2018).

*Flowering synchronization index*

For each of all the *H. halimifolium* plants within the population, we estimated the flowering phenology by registering the presence or absence of flowers every three days during the entire flowering peak season. To estimate individual overlap in flowering time we calculated a flowering synchronization level by modifying the index proposed by Marquis (1988) as follows:

$$S_i = \frac{\sum_{t=0}^{n} \left( \frac{x_t}{\sum_{t=0}^{n} x_t} \right) (p_t)}{n}$$

where $S_i$ is the mean flowering synchronization level of individual *i*, averaged over *n* censuses carried out during the peak flowering season, $x_t$ is the number of open flowers on individual *i* at time *t*, $\frac{x_t}{\sum_{t=0}^{n} x_t}$ is the proportion of open flowers on individual *i* of the total number of open flowers in that individual during the peak flowering season, *n* is the number of censuses during the peak flowering season, and $p_t$ is the proportion of individuals in flower at time *t* in the population.

*Female fitness estimation*

For each surveyed individual plant, the different components of the female plant fitness (i.e. fruit set, number of seeds per fruit and seed mass) were performed in inflorescences randomly selected and marked after performing the pollinator visitation surveys. An overall female plant reproductive success for each individual plant was estimated by weighing the total number of seeds per plant by seed mass as follows:

$$c_i = \frac{(\mathrm{SM}_i - \mathrm{SM}_{min})}{(\mathrm{SM}_{max} - \mathrm{SM}_{min})}$$

$$\mathrm{FRS}_i = \mathrm{NF}_i \times \mathrm{FS}_i \times \mathrm{NS}_i \times \mathrm{c}_i$$

where *SM* is the mean seed mass (mg), *NF* is the total number of flowers during the peak flowering season, *FS* is the fruit set (i.e. percentage of flowers setting fruit) and *NS* is the mean number of seeds per fruit.

*Topological position of individual plants*

Closeness centrality is positively related to the shortest number of direct and indirect interactions between one node and all other nodes in the network (Freeman 1978). It is based on the reachability of a node within a network, indicating how well pollen originating from a focal plant can reach other plants or how well pollen from conspecifics can reach this focal plant. Betweenness centrality reflects the proportion of shortest paths between two nodes going through the focal node (Freeman 1978), and therefore individual plants with high site betweenness act as bridges, connecting one part of a network to another (Newman 2004). It represents how well an individual plant facilitate the exchange of pollen among all nodes within the network. Centrality measures were estimated using the 'bipartite' R package (Dormann et al. 2008).

*Exponential random graph model (ERGM)*

Variables selected for analysis (here intrinsic and extrinsic plant attributes) can be combined into different ERGMs to identify the set of mechanisms that best explain the observed network. Hence, to determine which ecological variables are the main contributors to the network structure, we set four models including different predictors: full model, phenotypic traits model, microsite model excluding neighbourhood, and neighbourhood model. All predictor variables were previously scaled to allow meaningful comparisons among those variables. We calculated AIC (Akaike Information Criterion) weights for model selection (Link & Barker 2006) and performed goodness-of-fit diagnostics to assess the selected model fit to the data (Kolaczyk & Csárdi 2014).

**Table S1.** Classification of pollinator species visiting *Halimium halimifolium* plants into functional groups defined considering similarities in feeding habits, foraging behavior and body size.

| Pollinator functional group | Body length (mm) | Number of species | Taxa |
|---|---|---|---|
| Large bee | >20 | 3 | Hymenoptera: Apidae (*Bombus terrestris*, *Xylocopa violacea*, *Xylocopa cantabrita)* |
| Medium size bee | 10-20 | 4 | Hymenoptera: Anthophoridae (*Anthophora* sp.), Melittidae *(Dasypoda* sp.), Andrenidae (*Andrena* sp.), Apidae (*Apis mellifera)* |
| Small bee | <10 | 1 | Hymenoptera: Halictidae (*Lasioglossum* sp.) |
| Large beetle | >15 | 3 | Coleoptera: Alleculidae (*Heliotaurus ruficollis*), Meloidae (*Mylabris* sp.), Scarabaeidae (*Chasmatopterus illigeri)* |
| Medium size beetle | 5-15 | 5 | Coleoptera: Prionoceridae (*Lobonyx aeneus*), Nitidulidae, Oedemeridae |
| Small beetle | <5 | 3 | Coleoptera: Buprestidae (*Anthaxia dimidiata)*, Cantharidae *(Malthodes* sp.), Dermestidae (*Anthrenus* sp.) |
| Hoverfly | - | 2 | Diptera: Syrphidae (*Eristalis tenax*, *Episyrphus balteatus)* |
| Beefly | - | 1 | Diptera: Bombyliidae (*Phthiria* sp.) |

**Table S2.** Range, mean, median and standard deviation (SD) values of intrinsic and extrinsic attributes measured for individual *Halimium halimifolium* plants in the studied population.

| Plant attribute | Min | Max | Mean | Median | SD |
|---|---|---|---|---|---|
| ***Intrinsic*** | | | | | |
| Maximum height (m) | 0.185 | 1.926 | 0.844 | 0.752 | 0.416 |
| Flower guide size (mm) | 0.000 | 3.000 | 1.917 | 2.000 | 0.720 |
| Flower size (mm) | 25.138 | 37.540 | 31.789 | 32.124 | 2.767 |
| Total flower number | 3.000 | 5264.000 | 554.367 | 251.000 | 841.524 |
| Flowering synchrony | 6.821 | 67.949 | 18.675 | 16.126 | 10.786 |
| ***Extrinsic*** | | | | | |
| Distance to stream (m) | 66.025 | 78.069 | 75.537 | 71.562 | 25.432 |
| Distance to tree (m) | 0.000 | 40.360 | 16.674 | 15.534 | 10.261 |
| Distance to edge (m) | 72.872 | 78.305 | 41.747 | 75.891 | 12.900 |
| Intraspecific neighbours in r=1m (m2) | 0.253 | 25.231 | 6.573 | 5.031 | 5.478 |
| Interspecific neighbours in r=1m (m2) | 0.000 | 38.474 | 3.449 | 0.507 | 7.203 |
| Intraspecific neighbours in r=2m (m2) | 0.253 | 28.183 | 10.448 | 8.679 | 7.394 |
| Interspecific neighbours in r=2m (m2) | 0.000 | 95.728 | 7.325 | 1.233 | 15.674 |

**Table S3.** Standardized coefficients for the first canonical variable across original sets of variables, testing the overall correlation between visitation by pollinator functional groups and plant attributes including phenotype and microsite variables. Significance of the association between pollinator visitation to individual plants and plant attributes for the first canonical variable was tested with a Wilk's lambda test. High correlation coefficients (>0.2) appear in boldface.

| Original Variable | Canonical Variable I ($\lambda$= 0.047; R= 0.849***) |
|---|---|
| Beefly | 0.040 |
| Hoverfly | -0.137 |
| Large bee | **0.359** |
| Large beetle | **0.572** |
| Medium size bee | **0.591** |
| Medium size beetle | 0.153 |
| Small bee | -0.019 |
| Small beetle | 0.109 |
| Maximum height (m) | **-0.333** |
| Flower guide size (mm) | -0.006 |
| Flower size (mm) | 0.092 |
| Total number of flowers | **0.828** |
| Synchrony index | -0.012 |
| Distance to stream (m) | -0.155 |
| Distance to tree (m) | **0.223** |
| Distance to edge (m) | **-0.229** |
| Cover of intraspecific neighbours within 1m (m2) | **0.221** |
| Cover of interspecific neighbours within 1m (m2) | **-0.274** |
| Cover of intraspecific neighbours within 2m (m2) | **-0.397** |
| Cover of interspecific neighbours within 2m (m2) | **0.582** |
| | F= 1.67; df= 96, 279.72; p< 0.001 |

**Table S4.** Exponential random graph models (ERGMs) results with the structure of the bipartite pollination network as response variable. Four models with different predictors (full model, model with phenotypic traits, model with microsite characteristics excluding neighbourhood and model with neighbourhood variables) were performed and selected following AIC weights. Significant values ($p < 0.05$) appear in boldface.

| | Full model | | Phenotypic traits | | Microsite | | Neighbourhood | |
|---|---|---|---|---|---|---|---|---|
| | Estimate ± SE | z | Estimate ± SE | z | Estimate ± SE | z | Estimate ± SE | z |
| Edges | -0.834 ± 0.502 | -1.663 | 0.086 ± 0.241 | 0.357 | -0.472 ± 0.433 | -1.088 | 0.024 ± 0.225 | **0.107** |
| Individual plant | 0.003 ± 0.009 | 0.393 | -0.008 ± 0.006 | -1.329 | 0 ± 0.008 | 0.062 | -0.011 ± 0.006 | -1.909 |
| Maximum height (m) | 0.065 ± 0.126 | 0.515 | 0.318 ± 0.101 | **3.142** | | | | |
| Flower guide size | -0.089 ± 0.077 | -1.151 | -0.103 ± 0.072 | -1.424 | | | | |
| Flower size (mm) | -0.319 ± 0.086 | **-3.699** | -0.116 ± 0.063 | -1.839 | | | | |
| Total number of flowers | 0.25 ± 0.117 | **2.132** | 0.025 ± 0.103 | 0.246 | | | | |
| Flowering synchrony | -0.491 ± 0.148 | **-3.307** | -0.276 ± 0.095 | **-2.902** | | | | |
| Distance to stream (m) | -0.42 ± 0.194 | **-2.171** | | | -0.259 ± 0.169 | -1.536 | | |
| Distance to tree (m) | 0.035 ± 0.119 | 0.296 | | | -0.051 ± 0.086 | -0.596 | | |

| | | | | | | | | |
|---|---|---|---|---|---|---|---|---|
| Distance to edge (m) | 0.485 ± 0.174 | **2.798** | | | 0.226 ± 0.145 | 1.561 | | |
| Cover of intraspecific neighbours in r=1m (m2) | 0.03 ± 0.176 | 0.171 | | | | | 0.027 ± 0.123 | 0.224 |
| Cover of interspecific neighbours in r=1m (m2) | 0.554 ± 0.122 | **4.531** | | | | | 0.32 ± 0.096 | **3.322** |
| Cover of intraspecific neighbours in r=2m (m2) | 0.126 ± 0.168 | 0.750 | | | | | 0.133 ± 0.141 | 0.942 |
| Cover of interspecific neighbours in r=2m (m2) | -0.252 ± 0.157 | -1.604 | | | | | -0.029 ± 0.13 | -0.224 |
| AIC | 619.5 | | 650.8 | | 656.3 | | 646 | |
| AIC weights | 1 | | 0 | | 0 | | 0 | |

**Table S5.** Results of Moran's I analyses testing spatial autocorrelation in intrinsic attributes and female reproductive success (FRS) of individual plants. Observed Moran's I values close to -1 represent that the variable values are spatially dispersed while values close to 1 indicate a spatial clustering of similar values of the variable.

| Plant attribute | Observed Moran's I | Expected Moran's I ± SD | p |
|---|---|---|---|
| Maximum height (m) | 0.016 | -0.017 ± 0.026 | 0.205 |
| Mean height (m) | 0.029 | -0.017 ± 0.062 | 0.062 |
| Area (m2) | -0.018 | -0.017 ± 0.025 | 0.958 |
| Perimeter (m) | -0.016 | -0.017 ± 0.025 | 0.963 |
| Flower guide size (mm) | -0.030 | -0.017 ± 0.026 | 0.604 |
| Flower size (mm) | 0.005 | -0.017 ± 0.026 | 0.395 |
| Mean flower number | 0.007 | -0.017 ± 0.024 | 0.306 |
| Total flower number | -0.008 | -0.017 ± 0.022 | 0.673 |
| Flowering synchrony | -0.001 | -0.017 ± 0.024 | 0.450 |
| FRS | 0.006 | -0.017 ± 0.024 | 0.340 |

**Table S6.** Results of the relative importance of regressors analysis to assess the contribution of the different fitness components to the female reproductive success (FRS) of individual plants.

| | Estimate ± SE | t value | p | % Effect |
|---|---|---|---|---|
| (Intercept) | -0.111 ± 0.053 | -2.09 | **0.042** | |
| Total number of flowers | 0.0002 ± 0.000009 | 22.258 | **<0.001** | 0.861 |
| Fruit set | 0.219 ± 0.044 | 4.993 | **<0.001** | 0.040 |
| Number of seeds | 0.001 ± 0.001 | 0.701 | 0.486 | 0.007 |
| Seed mass | 44.437 ± 68.537 | 0.648 | 0.520 | 0.002 |

**Fig. S1.** Study population of *Halimium halimifolium* in Doñana Natural Park (37°07'52.2''N 6°31'40.9''W) **(a)** and different floral phenotypes of *H. halimifolium* plants found in the study population **(b-c)**.

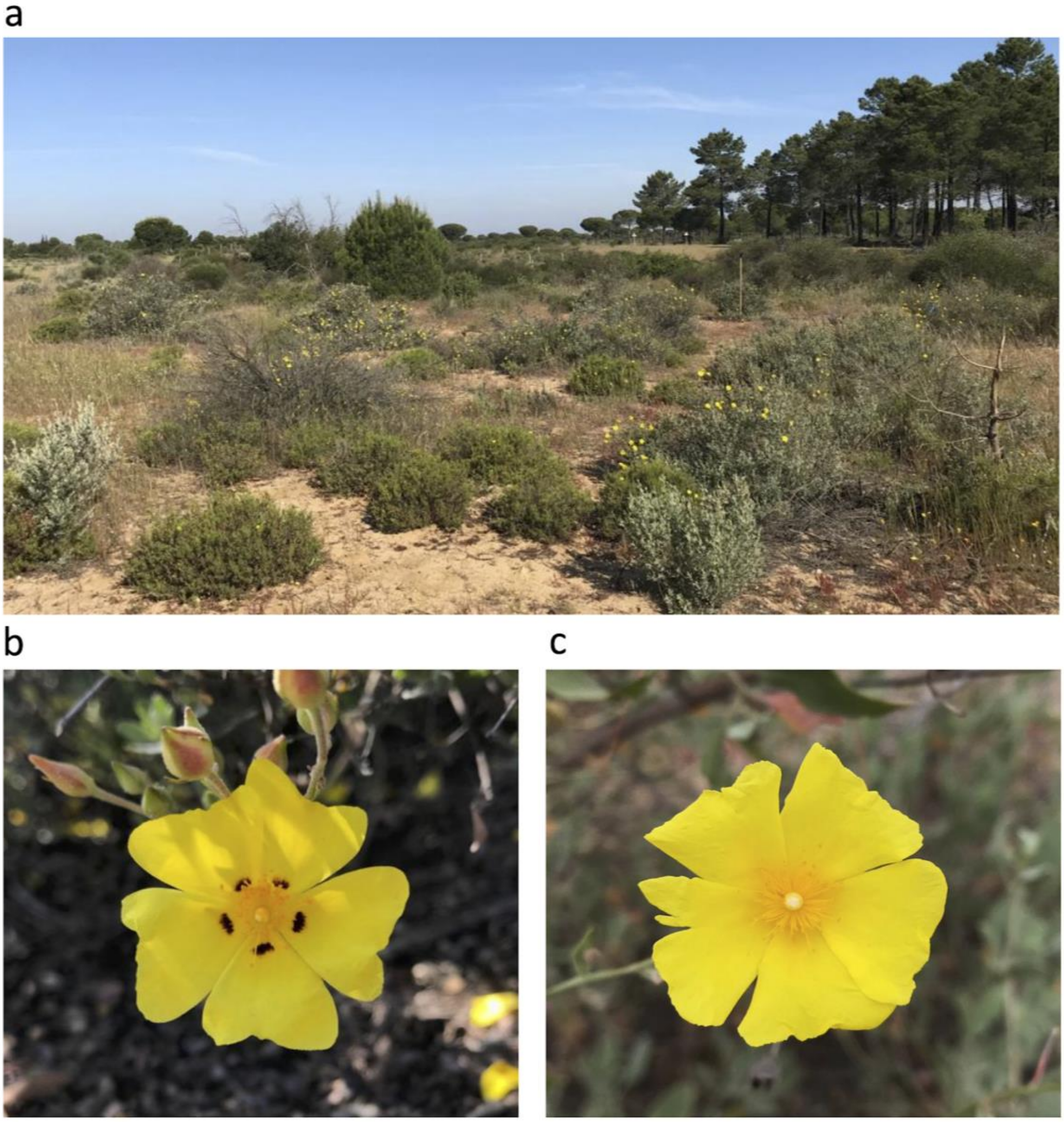

**Fig. S2.** Accumulation curves for the number of distinct pollinator functional groups **(a)** and pollinator species **(b)** recorded visiting the surveyed individual plants in the study population. The completeness of the survey was 100.00 ± 0.00 % (Chao asymptotic estimator ± SE) for pollinator functional groups and 82.35 ± 16.17 % for pollinator species. Boxplots indicate the median, interquartile range, range excluding outliers and outlier values for the estimated number of functional groups (a) or species (b) for increasing sampling effort; the shaded area shows the confidence interval for the resampled estimates.

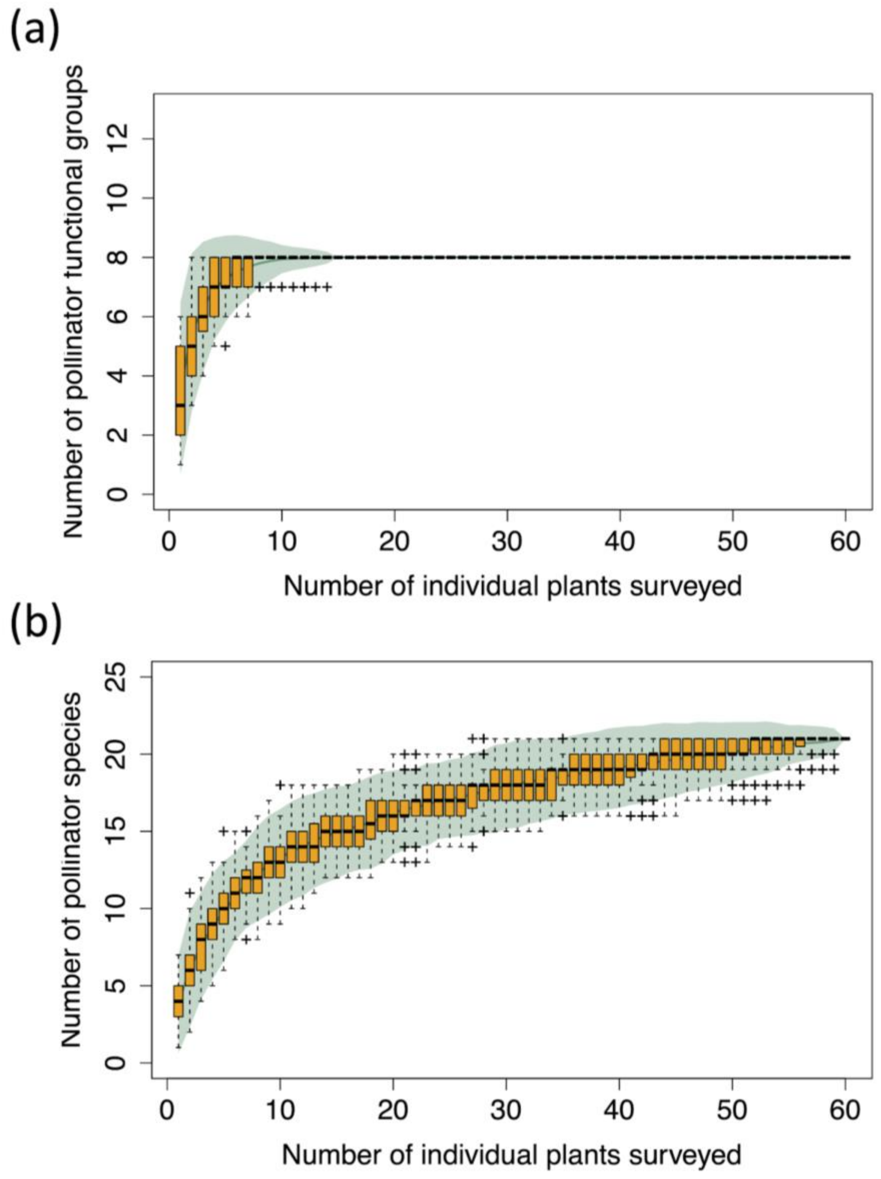

**Fig. S3.** Spatial distribution of all individual *Halimium halimifolium* plants surveyed in the studied population.

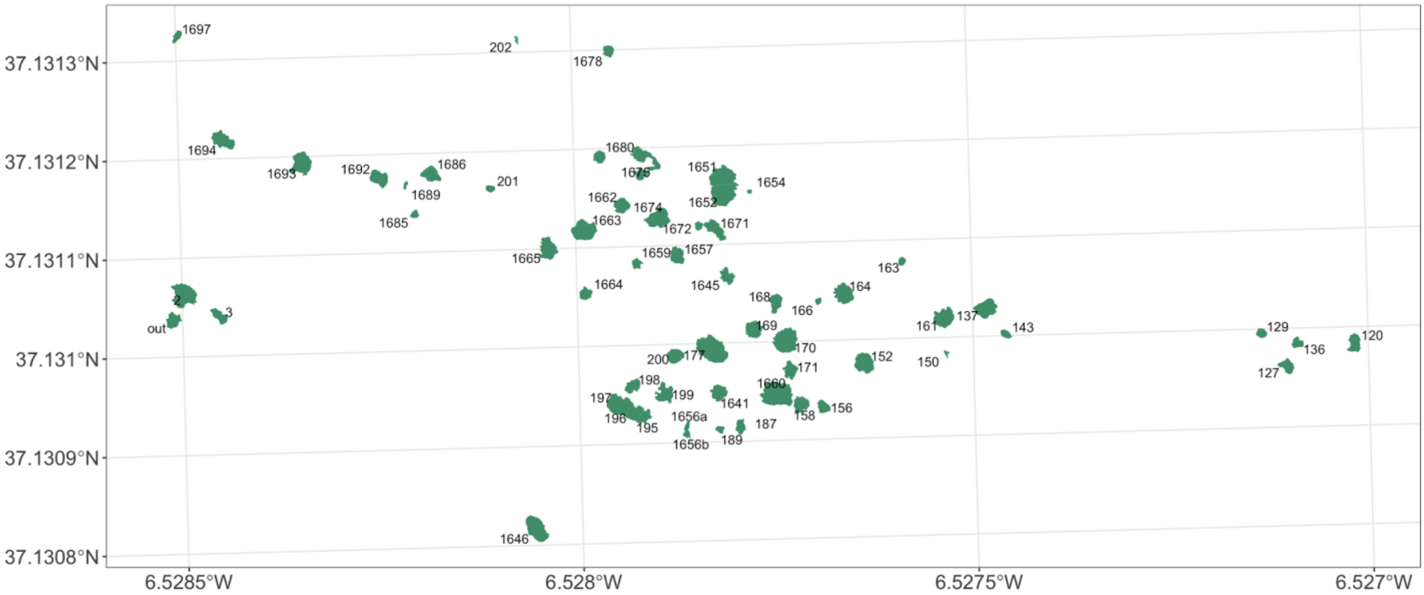

**Fig. S4.** Graph illustrating the weighted bipartite network between individual plants (green nodes) and pollinator functional groups (orange nodes). The links between nodes represent flower visitation interactions while the width of the links refers to the strength of the interaction (i.e. number of interactions recorded). Green node size is proportional to the female reproductive success (FRS) of individual plants. The layout of the network representation was created using an energy-minimization algorithm. Pollinator functional groups represent: Bf - Beefly, Hb - Hoverfly, Lbee - Large bee, Mbee - Medium size bee, Sbee - Small bee, Lbeet - Large beetle, Mbeet - Medium size beetle and Sbeet - Small beetle.

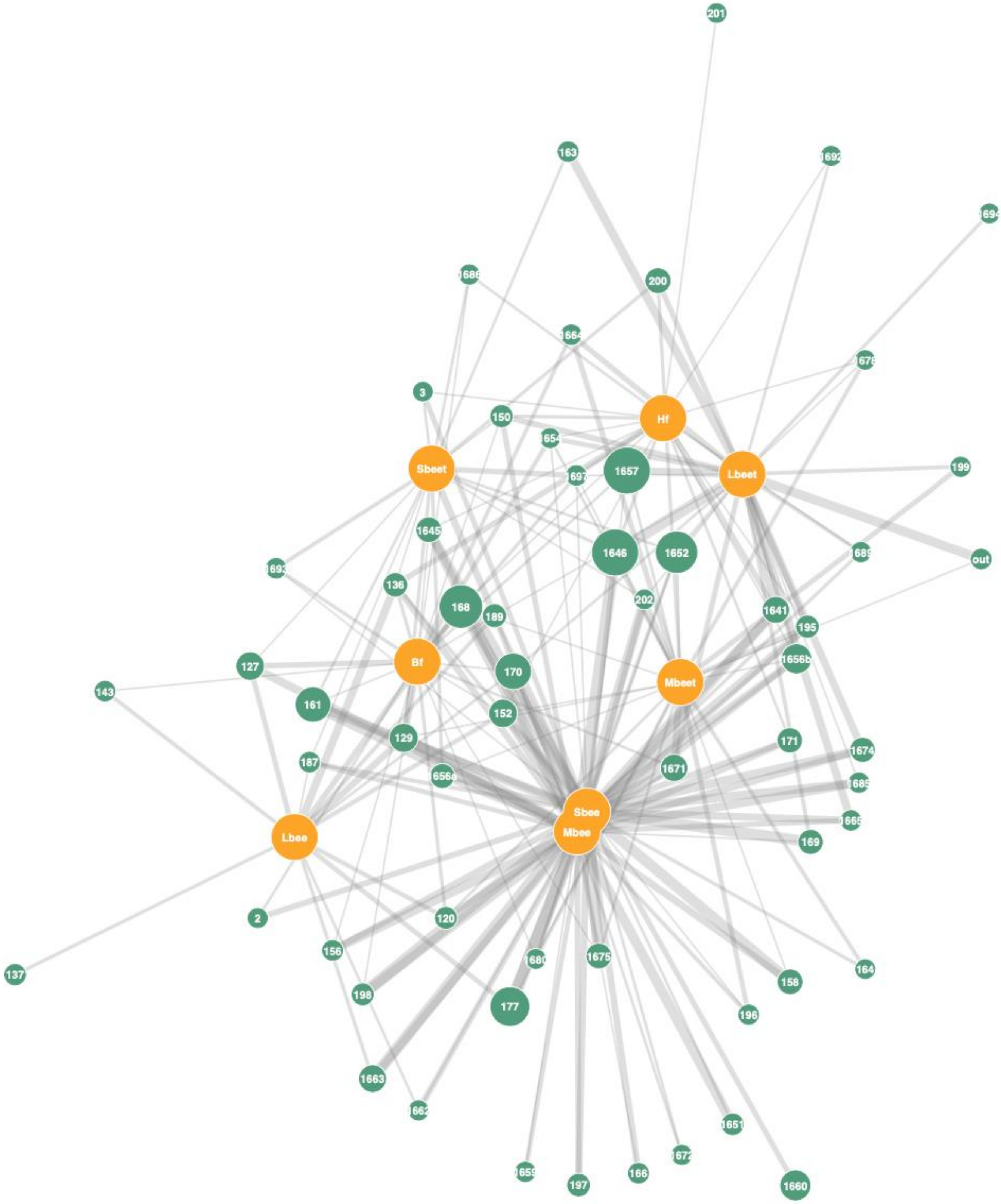

201
163
1692
1694
1686
200
1664
1678
3
150
Hf
1654
Sbeet
1697
1657
Lbeet
199
1645
1646
1652
1693
out
1689
136
168
189
202
1641
195
127
Bf
170
1656b
Mbeet
143
161
152
129
171
1674
187
1656a
1671
1685
Sbee
Mbee
1665
Lbee
169
2
120
156
1680
1675
164
158
198
177
196
1663
1662
1651
1672
1659
197
166
1660
